# The scent of symbiosis: gut bacteria affect social interactions in leafcutting ants

**DOI:** 10.1101/335521

**Authors:** Serafino Teseo, Jelle S. van Zweden, Luigi Pontieri, Pepijn W. Kooij, Søren J. Sørensen, Tom Wenseleers, Michael Poulsen, Jacobus J. Boomsma, Panagiotis Sapountzis

## Abstract

Animal gut microbiota affect host physiology and behaviour. In eusocial Hymenoptera, where colony-level integrity is preserved via a nestmate discrimination system based on cuticular hydrocarbon mixtures, microorganismal effects may influence social dynamics. Although nestmate recognition has undergone a thorough exploration during the last four decades, few studies have investigated the putative role of gut microbes. Here we integrate metagenomic, chemical and behavioural approaches to test whether gut microbes affect nestmate recognition in *Acromyrmex echinatior* leaf-cutting ants. Treating workers with a sterile diet or with antibiotics resulted in a substantial alteration of their gut microbial communities. In pairwise social interactions, untreated vs. antibiotic-treated nestmates behaved more aggressively than other nestmate and non-nestmate pairs, suggesting that the suppression of microbes indirectly alters chemical social cues and triggers aggressive behaviour. Chemical analyses on treated individuals revealed a decrease in the abundance of two metapleural gland antifungal compounds, and we confirmed the correspondence between aggression levels and chemical profile differences. Feeding microbiota-remodelled ants with conspecific faecal droplets partially restored the original bacterial communities. Furthermore, non-nestmates fed with faecal droplets from different colonies were unusually aggressive compared to pairs fed with faecal droplets from the same colony. This suggests that chemicals derived from microbial strains may shape nestmate recognition, opening novel questions about the role of microorganisms in the evolution of social behaviour.

## Introduction

In the evolution of mutualistic relationships between metazoans and prokaryotes, animals have co-opted the metabolic versatility of microbes to upgrade their physiology, while microorganisms have found favourable environments in animal bodies. As part of the physiology of their animal hosts, symbiotic microbes are also involved in their behavioural processes (1–5). Many insightful discoveries about the physiological and behavioural effects of symbiotic microorganisms stem from the study of germ-free or germ-remodelled animals. Research comparing germ-free mice to their untreated counterparts has revealed microbial gut symbionts to affect anxiety-like behaviour (6–8) and social interactions (9). Similarly, a flourishing corpus of *Drosophila* studies suggests that gut microbes mediate a plethora of physiological/behavioural processes, including specific appetites for proteins (10), mate choice and mating dynamics (11–14) and the recognition of kin and familiar individuals (15).

This increasing awareness about the behavioural role of microbes opens questions about how microorganisms may affect the behavioural ecology of animals with radical social adaptations, such as the eusocial insects. These live in family groups with a permanent division of reproductive labour, worker castes and sophisticated chemical communication (16). Eusocial insect colonies exhibit complex behaviours resulting from the cooperative interactions of individuals, such as foraging, nursing and nest construction, as well as policing and colony defence against unrelated intruders. Across taxa, group-level integrity is ensured by a chemical-based nestmate recognition system, which relies on signals encoded in long-chain hydrocarbons forming a waxy layer on the insect cuticle (cuticular hydrocarbons, or CHC).

Our current knowledge about the microbial effects on eusocial insect CHC-mediated social behaviour is limited and inconsistent. *Reticulitermes speratus* termites fed with bacteria extracted from individuals of unrelated colonies are attacked by nestmates, and antibiotic-mediated manipulation of bacterial communities affects nestmate recognition behaviour (17). In *Pogonomyrmex barbatus* harvester ants, individuals with experimentally-augmented cuticular microbiomes are rejected by nestmates more than controls, whereas antibiotic-treated individuals are not. This suggests that cuticle-dwelling microbes influence nestmate recognition dynamics (18). Contrarily, however, topical antibiotic administration on *Acromyrmex subterraneus* leafcutter ants does not affect cuticular hydrocarbon profiles (19). Similarly, a study on *Camponotus* carpenter ants revealed a negative correlation between the levels of the bacterial gut symbiont *Blochmannia* and CHC quantities, whereas relative CHC proportions were not affected (20). Finally, antibiotic administration affects interspecific but not intraspecific social interactions in the Argentine ant *Linepithema humile*, suggesting gut microbiota not to be involved in nestmate recognition in this ant species (21).

Despites providing insights into the role of symbiotic microbes in CHC-mediated nestmate recognition, these studies either investigate microbial effects through behavioural tests, without correlating CHC measures (17,21), or compare CHC profiles between antibiotic-treated and control individuals without behavioural tests (18–20). To gain more insights into the interplay between gut microbiota and chemical-based social interactions, we here seek to implement an integrative analysis of CHC, gut bacterial communities and nestmate recognition behaviour. We remodelled the previously-characterized gut microbiota of *Acromyrmex echinatior* leafcutter ants (22,23) to investigate the link between gut bacterial communities and socially relevant chemical signals. Our hypothesis was that remodelling of the ant gut microbiota would result in a chemical profile shift with measurable effects on social interactions. Therefore, in a first set of experiments, we either mildly suppressed or completely eliminated the native gut microbiota of *A. echinatior* using respectively a sterile sucrose diet or antibiotics (Round 1). We then characterized the effects on the ant gut microbial communities and chemical profiles, and used dyadic aggression trials to evaluate treatment-dependent changes of gut microbial communities with chemical profiles and behaviour. In a second set of experiments, we partially restored the original ant gut microbial communities with the aim to rescue the chemical and behavioural effects obtained with the experimental diet and the antibiotics (Round 2). To achieve this, we fed experimentally-treated individuals with faecal droplets of untreated conspecifics and again analysed their microbial communities, chemical profiles and social interactions.

## Material and methods

### Experimental design

We used workers from four *A. echinatior* colonies (Ae150, Ae322, Ae153 and Ae331, hereafter named respectively A, B, C and D) with an already characterized gut microbiota (24,25) for the experiments (Figure 1). Colonies were collected in Gamboa, Republic of Panama in 2003-2010, and kept at 25°C, 70% RH and 12:12 L:D photoperiod. Individuals were taken from fungus gardens (375 individuals/colony, N=1500) and placed in groups of 15 in sterile Petri dishes (ø 90×15mm; 25 dishes/colony, N=100) including a food container. For each colony, we randomly assigned workers to two treatment groups: 1) ants kept on a sterile 10% sucrose solution diet (150 individuals/colony); 2) ants kept on the same sterile 10% sucrose solution supplemented with 1mg/ml of the antibiotic tetracycline (225 individuals/colony). After two weeks, 522 individuals were tested in a first series of experiments (Round 1) including aggression assays and analyses of CHC (GC-MS, see below) and gut microbiota (qPCR and MiSeq, see below). The remaining individuals underwent the experiments in Round 2, where we attempted to restore their original microbiota by feeding ants on a sterile 10% sucrose solution supplemented with faecal droplets (0.033 droplets/μl) obtained by squeezing abdomens of untreated workers from the source colonies. Colony B and D workers received nestmate droplets, whereas A and C workers received non-nestmate droplets (from colonies B and D, respectively, Figure 1). After one week, individuals were tested in aggression assays and used for CHC and microbiota analyses. Throughout the experiments, we monitored survival to determine the effects of our treatments on ant mortality (details in the Supplementary information, Figure S1).

**Figure 1.**
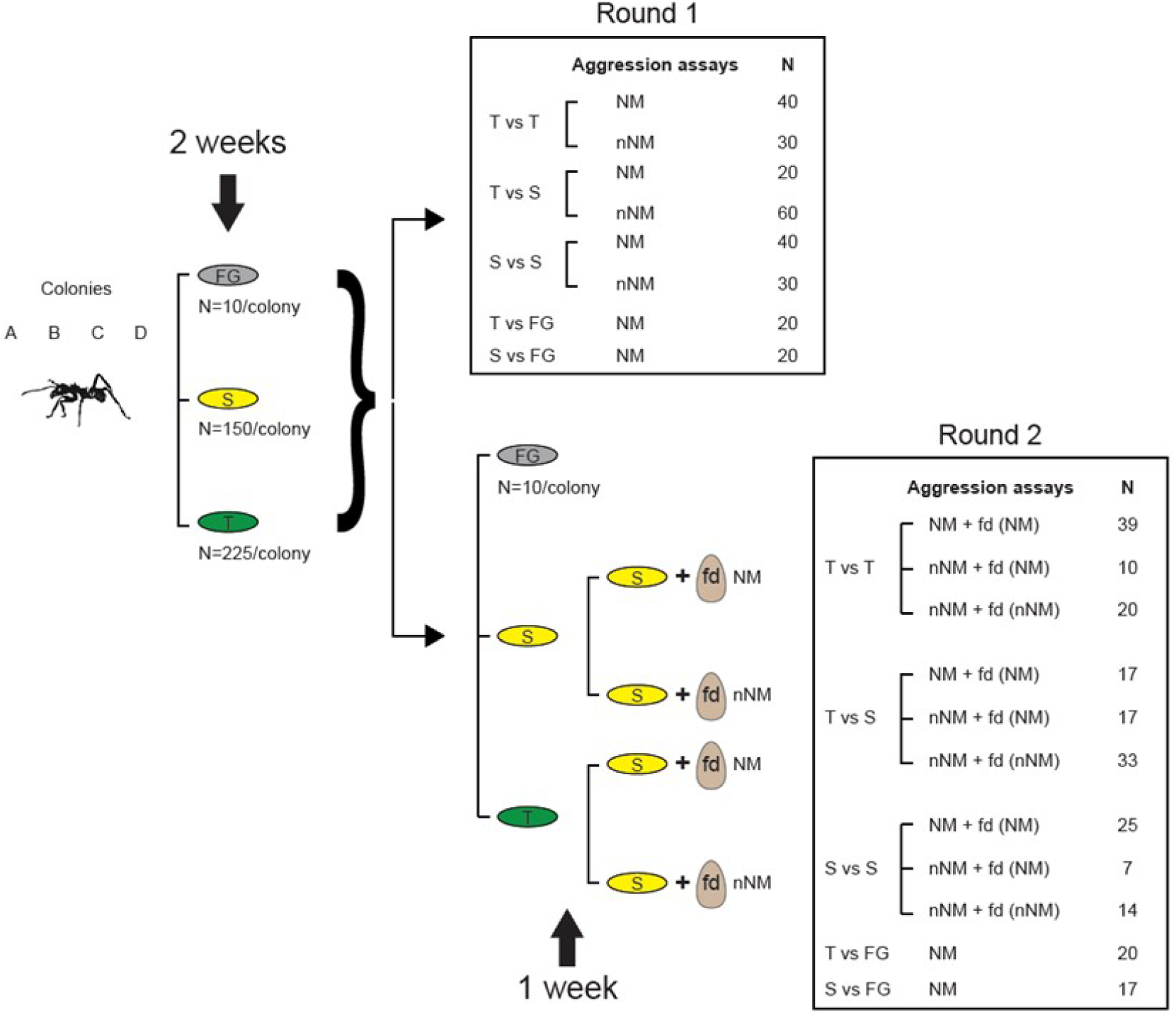
Experimental design. Ants were collected from fungus gardens (FG) and kept for two weeks on a sucrose (S) or Tetracycline (T) diet (Round 1). After two weeks we performed aggression trials among the three treatment groups. Individual guts were then dissected for gut bacterial characterization (16S-qPCR and 16S-MiSeq), whereas thoraxes and heads were used for chemical extractions and subsequent GC-MS analyses. In Round 2, the remaining individuals were fed for one week on faecal droplets (fd) either from their original nestmates (NM) or non-nestmates (nNM) from a different colony. These ants were then tested in aggression trials similar to Round 1 and collected for gut microbiota characterization and GC-MS analyses.

Finally, we set up an independent smaller-scale experiment in which we compared the microbial communities of ant guts, heads and thoraxes before and after the experimental diet treatment (details in the Supplementary Information, Figure S2).

### Aggression assays

We tested microbiota-remodelled and untreated ants against nestmates and non-nestmates from the original colonies, in both experimental rounds (3-5 replicates/combination, total=260 tests; Figure 1, Data set S1). Assays consisted of dyadic encounters in Petri dishes with clean filter paper on the bottom. For two minutes after first contact, an observer (blind with respect to ant treatment and original colony) used the software Etholog 2.25 (26) to quantify the frequency and duration of biting, mandible opening, antennation and absence of contact (Figure S3). For statistical analyses, we excluded the mandible opening behaviour (usually considered as aggressive) because pooling it with biting produced similar results (Supplementary Information). In addition, because keeping ants in the same Petri dishes for three weeks resulted in a significant effect on aggression (Binomial GLMM with ‘Petri dish’ as fixed and ‘colony’ as random variables, z=3.21, Df=141, p=0.001), we considered only interactions between nestmates kept in different Petri dishes.

We analysed data in R using the packages *lme4, car* and *multcomp* (27–29), fitting generalized linear mixed models (GLMMs). Classifying biting as aggressive behaviour and antennal contact as non-aggressive allowed measures to be analysed as a single binomial response variable. For Round 1, the initial model included biting/antennation frequencies as response variable, whereas nestmate, diet treatment and their interaction were included as fixed factors. The colony origin of experimental individuals was included as random effect term. The model used for Round 2 included the same response variable as Round 1 model, and ‘faecal droplets’ was included as an additional fixed factor (two levels: from nestmates or non-nestmates). We tested the significance of fixed effects using the *car* function ‘Anova’. Where needed, we conducted post hoc planned contrasts between groups of interest using the *multcomp* function ‘glht’, correcting alpha values with false discovery rate (FDR).

### DNA extractions

Four to eight workers per treatment per colony used in the aggression assays were ice anesthetized and individually dissected in sterile Phosphate Buffered Saline (PBS). Pooled crop, midgut, hindgut, Malpighian tubules and fat body cells were stored at −20°C until DNA extractions, and immediately homogenized after thawing in 200μl ATL buffer supplemented with 20μl proteinase K (Qiagen) using sterile pestles. Subsequently, ø0.45 mm glass beads were added and tubes vortexed for 30s, after which the samples were incubated at 56°C overnight under constant agitation. DNA was extracted using the Qiagen Blood and Tissue kit, and all samples were eluted in 100μl AE elution buffer.

### 16S rRNA qPCR analyses

The gut microbiota of *A. echinatior* normally consists of five predominant OTUs (Operational Taxonomic Units, representing a cluster of bacterial 16S rRNA sequences of ≥97% similarity) (22,25,30). These belong to the genus *Wolbachia* (*wolAcro1,* including two strains: *w*SinvictaA and *w*SinvictaB) and the orders Entomoplasmatales (class: Mollicutes, OTUs *EntAcro1, EntAcro2* and *EntAcro10*) and Rhizobiales (class: Alpha-Proteobacteria, OTU *RhiAcro1* (25,31–33)). In order to monitor how the dietary treatment affected the communities, we screened individual worker guts with qPCR (detailed methods in Supplementary Information, text and Table S1) on the five most abundant OTUs (Data set S2, procedures described in (25)), with Cycle threshold (Ct) mean of replicated samples used as a measure of amplicon abundance. The elongation factor 1 alpha (EF-1α) was used as a reference gene (34). Each run included two negative controls with no added template for each gene used. Data were ordinated using an unscaled principal coordinate analysis (PCoA) and inter-sample distances were again calculated using Canberra, Hellinger and Bray-Curtis methods. We estimated the difference in variation among groups using the HOMOVA command (with Bonferroni-adjusted alpha-values) implemented in mothur (35). For analysis, we initially used a standard curve with PCR products in tenfold dilution series of known concentration (fold change method) to calculate the PCR efficiency using the REST software (36). Data were imported in R and expressed as ΔΔC_T_ values, i.e. as the fold change relative to the EF-1α control gene (37), always using zero as reference. We used linear mixed models (LMM) with the ΔΔC_T_ values as response variable, ‘diet treatment’ (i.e., untreated, sugar-treated or tetracycline-treated) and ‘experimental round’ (i.e. Round 1 or 2) as fixed variables, and ‘colony’ (A, B, C or D) as random variable. Tukey post hoc tests were performed to evaluate significant differences between groups. We used GLMs to pair aggression data with distances calculated using ΔC_T_ values fold change differences, or with absolute copy numbers calculated using the qPCR data. All correlations were performed in R using the lme4, vegan, effects, Rmisc and ggplot2 packages (27,38–41).

### 16S rRNA MiSeq analyses

To investigate whether novel OTUs appeared with the dietary treatment, we screened ant worker guts individually with 16S rRNA MiSeq sequencing. Amplicons were generated using the 515F/806R primers targeting the 16S rDNA V4 region (42), purified using the Agencourt AMPure XP (Beckman Coulter) and quantified using Quant-iT dsDNA High-Sensitivity Assay Kit and Qubit fluorometer (Invitrogen) to allow for dilution and mixing in equal concentrations before sequencing. Sequencing (for details see the Supplementary Information) took place in the Section of Microbiology at the University of Copenhagen using an Illumina MiSeq. Data [Genbank: SAMN04261407 - SAMN04261536 and SAMN05362797 - SAMN05362832] were analysed using mothur (35). Details on the mothur procedure are given in the Supplementary Information. After filtering/processing the sequencing data and clustering at 97%, rarefaction tables were constructed using pseudo-replicate OTU datasets containing 1-272000 sequences with 1000 iterations/pseudo-replicate, and the resulting curves were visualized in Microsoft Excel 2013. The final OTU Table was rarefied at 5000 reads after manual inspection of the rarefaction curves, which reduced the number of OTUs to 1500. We used the MiSeq data to calculate Canberra and Bray-Curtis distances, after which we used Non-metric MultiDimensional Scaling (NMDS) to ordinate and visualize the effects. We used GLMs to pair aggression data with Bray-Curtis or Canberra distances calculated from the 16S rRNA MiSeq data. Correlations were performed in R v3.2.3 using the lme4, vegan, effects, Rmisc and ggplot2 packages (27,38–41).

### Cuticular hydrocarbon analyses

CHCs were extracted by immersing the dissected heads and thoraces of aggression test individuals first in 150 μl HPLC-grade hexane, and then in 150 μl HPLC-grade chloroform (chemicals from Sigma-Aldrich, Belgium), both for 10 min under continuous agitation. The two extracts were mixed, and the solvent evaporated at room temperature in a laminar flow cupboard. The dry extract was then dissolved in 30μl hexane, of which 3μl were injected in a Shimadzu QP2010 Ultra GC-MS (splitless injector mode). Details on the GC-MS settings are given in the Supplementary Information. Our initial integration analysis of GC-MS runs detected 137 peaks, of which we selected 73 that had a relative abundance larger than 0.1% (Data set S3). Peak areas of cuticular compounds were integrated using R v3.1.0 (using package *xcms*, script available upon request) and normalized using a Z-transformation (43). To compare odour profiles among different rounds, we used linear mixed models (LMM) with the relative abundance of each compound as the response variable, ‘diet treatment’ and ‘experimental round’ as fixed variables, and ‘colony’ as a random variable. We conducted linear hypotheses using the *multcomp* R package (44) function glht to evaluate differences between diet treatments in the same Rounds, and between the same diet treatments across Rounds. All p-values were corrected for false discovery rate (FDR) given that we had conducted 73 separate tests per contrast. We used a coinertia analysis (45) to check for correlations between CHCs (transformed to logarithmic data) and qPCR measures (ΔCt values) of the six most abundant bacterial taxa (see details about these taxa below). In short, we generated two independent data matrices (either using the individual profiles or the pairwise differences for each trial), performed PCA analyses and paired them using the coinertia analysis using a Monte-Carlo test with 10000 permutations.

## Results

### Survival analysis

During Round 1, mortality increased in tetracycline treated workers (Cox proportional hazard model, p < 0.001), similar to what had been observed in a previous study (25). However, this effect disappeared in Round 2, when all ants were fed on faecal droplets (Figure S1), suggesting that the harmful effect of tetracycline lasted only as long as it was administered to the ants.

### Aggression tests (Round 1)

In Round 1, where ants were isolated from their original colonies and fed on sterile sucrose diets with/without antibiotics (Figure 1), we found a significant nestmate*treatment interaction (χ^2^=6.9803, df=2, p=0.0305). The diet treatment had a significant effect on aggression, whereas being non-nestmates did not (diet treatment: χ^2^=29.62, df=4, p<0.001; nestmates vs. non-nestmates: χ^2^=1.64, df=1, p=0.18; N=260 dyadic aggression assays). This result was mostly due to the high biting frequency between tetracycline-treated ants and their untreated former nestmates taken from fungus gardens, which was significantly higher from all other nestmate trials (all p<0.05, Figure 2A, Table S2). For tetracycline-treated and sucrose-treated ants, aggression levels between non-nestmates were not significantly different from those observed in nestmate trials (tetracycline-treated: z=0.365, p=0.715, Figure 2A; sucrose-treated: z=1.284, p=0.287). Contrarily, in the sucrose-vs. tetracycline-treated groups, non-nestmate trials showed higher aggression than nestmate trials (z=2.375, p=0.045).

**Figure 2.**
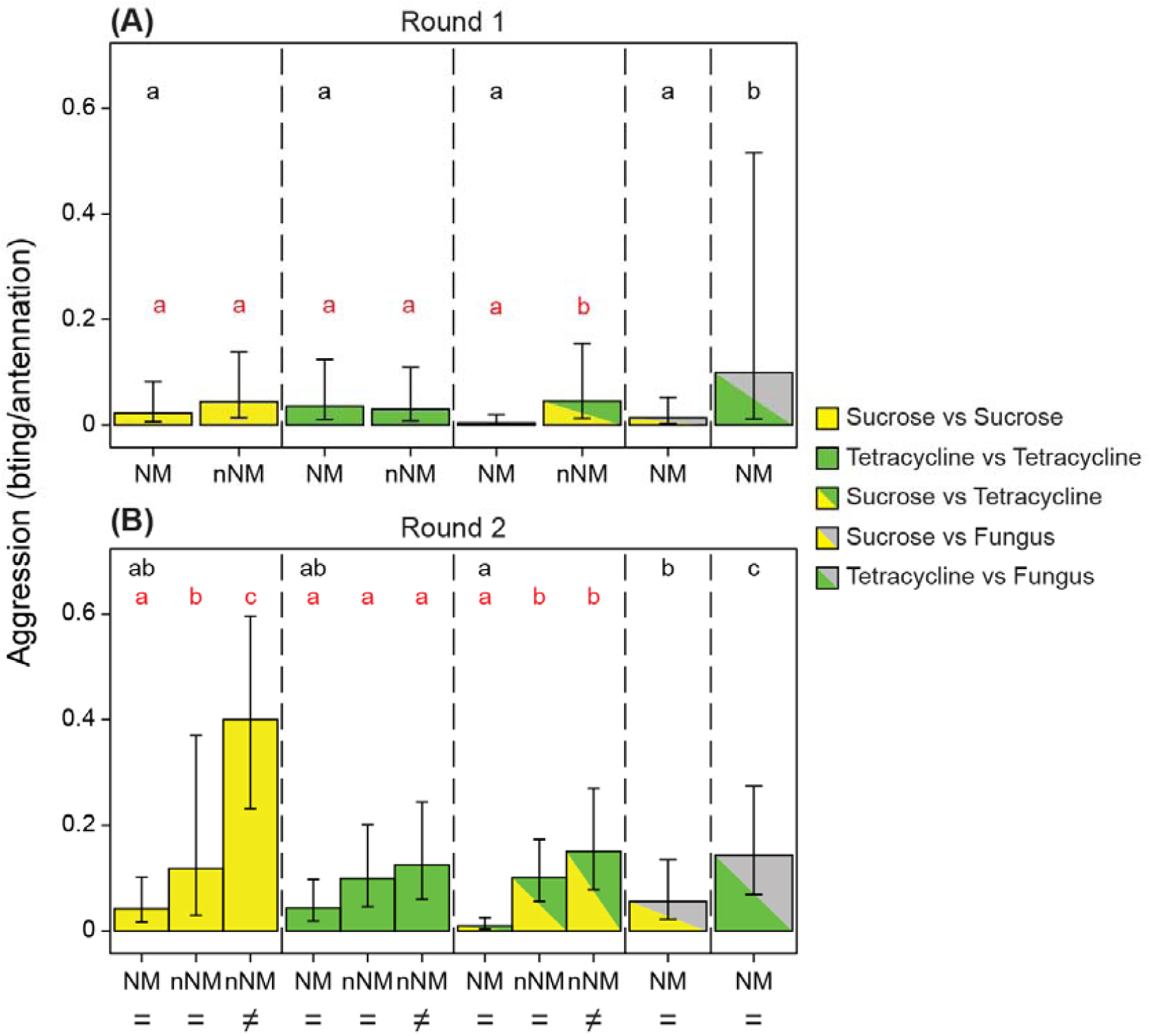
Effects of sucrose and tetracycline treatment on aggression. Mean biting/antennation ratio, with upper and lower confidence limits as error bars. **A.** Round 1. Tetracycline-treated vs. untreated nestmate ant pairs show the highest aggression levels, surpassing non-nestmate trials. **B.** Round 2. Encounters between individuals fed on faecal droplets. In nestmate trials, tetracycline-treated vs. untreated individuals still show the highest aggression. Letters indicate significance in pairwise comparisons (different letters = statistically significant differences). Red font indicates comparisons within treatment category (e.g. sucrose vs sucrose, tetracycline vs tetracycline) whereas black font shows comparisons across treatment categories (e.g. trials between sucrose-treated nestmates compared to trials between tetracycline-treated nestmates). In non-nestmate encounters, sucrose-treated individuals fed with faecal droplets from different colonies (≠) show the highest aggression levels, whereas non-nestmates fed with faecal droplets from the same colonies (=) exhibit low aggression. NM = nestmates; nNM = non-nestmates; FD= Faecal droplets. See Table S2 for details on statistical comparisons among treatment groups.

### Aggression tests (Round 2)

During Round 2, across sucrose-treated pairs, tetracycline-treated pairs and sucrose-vs tetracycline treated pairs, aggression was low in nestmate encounters fed with nestmate faecal droplets, higher in non-nestmate encounters fed with nestmate faecal droplets and maximal in non-nestmate encounters fed with non-nestmate faecal droplets (Figure 2b). In particular, the highest aggression levels appeared in encounters between non-nestmate sucrose-treated ants fed with faecal droplets from different colonies (z=2.05, df=85, p=0.041; Figure 2b, Table S2). Tetracycline-treated pairs showed instead low aggression (Figure 2b), regardless of whether they were fed on the same or different faecal droplets. Sucrose-treated vs tetracycline-treated ants exhibited low aggression levels, similar to tetracycline-treated pairs. We found a significant effect of the interaction faecal droplet*treatment (χ^2^=7.57, df=2, p<0.05), but not diet treatment*nestmate (χ^2^=3.36, df=2, p=0.18; Figure 2b, Table S2). All three main effects were significant (faecal droplets: χ^2^=7.99, df=1, p<0.05; diet treatment: χ^2^=36.97 df=4, p<0.001; Nestmate: χ^2^=5.2437 df=1, p<0.05).

### Gut bacterial communities changes (Round 1)

We examined the changes of the gut bacterial communities using both 16S rRNA MiSeq amplicon sequencing and 16S rRNA qPCR to measure the levels of the five most abundant OTUs (Data set S2; (22,25,30). Microbiomes of tetracycline-treated individuals were most strongly affected, showing the lowest variance, whereas untreated individuals collected from fungus gardens showed the highest variance (HOMOVA, p<0.001; Figure S4). While gut bacterial communities of ants reared on fungus gardens or sucrose differed significantly from those of tetracycline-reared ants (p<0.001 and p=0.013, respectively; Figure S4), their communities were different, even though not significantly, from each other (p=0.053). Furthermore, diet had the strongest effect on the differences between bacterial communities of treatment groups (PERMANOVA, F_2,53_=9.162, p<0.001), while colony origin did not (PERMANOVA, F_3,53_=1.758, p=0.931). When examining each of the abundant OTUs individually, *EntAcro1* and *RhiAcro1* decreased in tetracycline- and moderately in sucrose-treated individuals, while *w*SinvictaB was largely unaffected by diet (Figures 3, S5a). *w*SinvictaA (present only in colony C, Data set S2), *EntAcro2* and *EntAcro10* (present respectively in 9 and 20 of 37 tested untreated workers taken from fungus gardens; Data set S2), increased slightly in sucrose- and decreased in tetracycline-treated ants, but these effects were not significant. The head, thorax and gut microbial community comparisons showed that the most marked change were of the gut bacterial communities (detailed results in the Supplementary Information; Figure S5).

**Figure 3.**
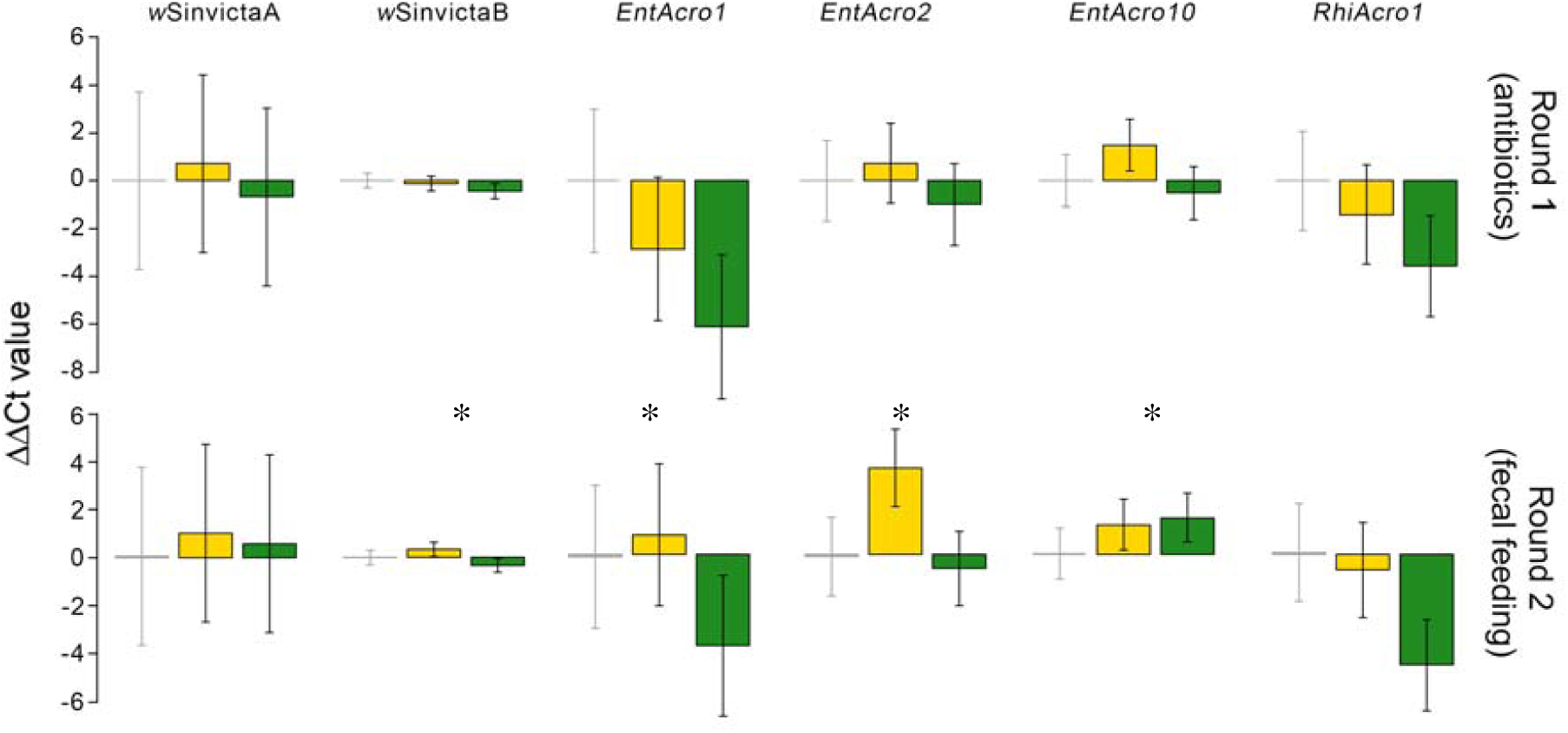
A. qPCR bacterial titres fold change of the six most abundant bacteria after experimental Rounds 1 and 2. Fold change is normalized to ant EF-1α copies, and the microbial strain levels of untreated conspecifics (ants reared on FG) are used as reference level (y=0). Scale bars represent standard errors. Asterisks indicate significance levels, p<0.05. Yellow: sucrose-treated; green bars: tetracycline-reared ants.

### Gut bacterial communities changes (Round 2)

The NMDS ordination showed that the gut bacterial communities of tetracycline-treated individuals before and after faecal droplet feeding were not significantly different. Contrarily, the gut bacterial communities of sugar-treated individuals were closer to those of untreated workers (from original fungus gardens) after faecal droplet feeding (Round 1), suggesting a shift towards the original communities (Figure S5). Interestingly, the gut bacterial samples of sugar-treated individuals exhibited a clear separation depending on which faecal droplets (nestmates or non-nestmates) they were fed on (PERMANOVA, F_1,43_=23.17, p<0.001 and F_1,43_=12.10, p<0.001, respectively and F_1,43_=5.04, p=0.004 for their interaction), whereas there was no such effect in the tetracycline-treated group (Figure S5). Tetracycline-treated ants undergoing faecal droplet feeding showed an increase of all OTUs but *EntAcro2* and *RhiAcro1*, whose levels were further reduced; sucrose-treated ants showed an increase of all gut bacterial taxa examined (Figure 3), but only changes in *EntAcro1, EntAcro2, EntAcro10* and *w*SinvictaB (respectively: t=-2.69, p=0.008; t=-2.02, p=0.044; t=-5.32, p<0.001; t=-5.95, p<0.001) were significant.

### Regression of bacterial changes and aggression

To identify which of the OTUs best explains aggression, and since the high variation both in aggression and gut microbiota composition precluded us from comparing mean effects, we regressed the observed aggression between nestmate test pairs on differences in abundance of their gut bacteria (Figure 4). Foremost of all, this analysis confirmed that more aggression was observed when nestmate pairs differed more in their gut microbial community (1500 MiSeq OTUs, using Bray-Curtis distances between pairs, binomial GLMM with ‘distance’ and ‘round’ as fixed variables and ‘colony’ as random variable, z=4.91, p<0.001). For individual taxa we used the more accurate qPCR data focusing on the main five taxa, regressing the observed aggression against their ΔΔCt values for each of the six gut bacterial OTUs (Figure 4). This analysis showed that aggression was not positively affected by either of the two *Wolbachia* strains (Figure 4; *wSinvictaA*: z=-0.66, p=0.509; *wSinvictaB*: z=-3.75, p<0.001), or *EntAcro2* (z=-0.872, p=0.383), but showed significant positive effects for *EntAcro1* (z=2.10, p=0.035), *EntAcro10* (z=6.09, p<0.001) and *RhiAcro1* (z=5.85, p<0.001). However, only differences in abundance of *RhiAcro1* had a strong effect on aggression in both rounds of the experiment (Figure 4; Data set S4; Round 1: z=3.25, p<0.001; Round 2: z=4.17, p<0.001), suggesting that Rhizobiales are a driving force in the recognition of nestmates (see Data set S4; similar results were obtained when both biting and mandible opening were treated as aggressive behaviors).

**Figure 4.**
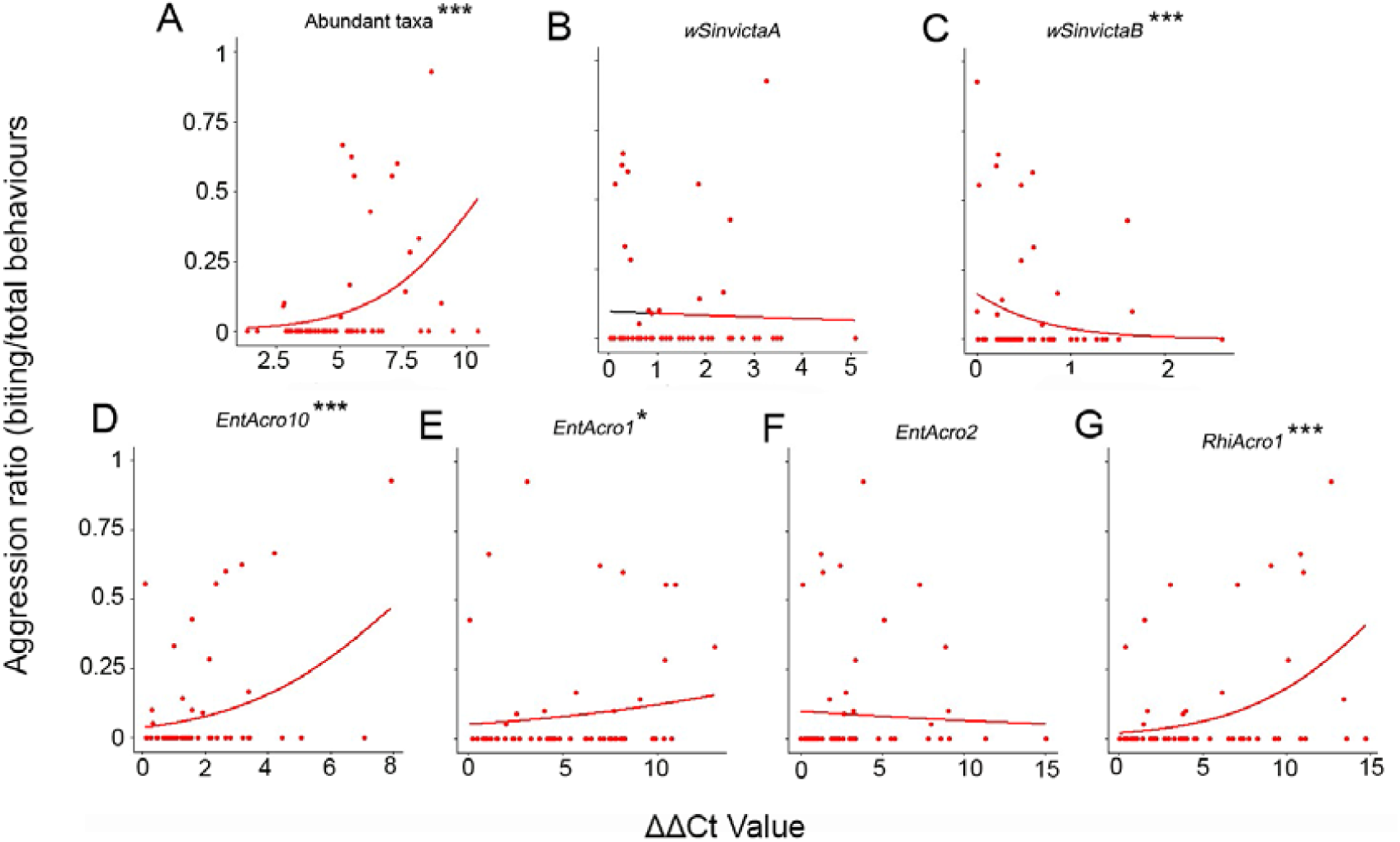
A. Association between aggression level and differences in the abundant gut bacteria using GLMMs. The relationship between biting frequency and bacterial abundances measured using the qPCR fold change (Ct distance) for both rounds for all six or for individual bacterial taxa. Curves represent predicted values from the models. Asterisks indicate significance levels: * p<0.05, ** p<0.01, and *** p<0.001.

### Effects of diet on CHC profiles

Compared to untreated individuals from original fungus gardens, sucrose- and tetracycline-treated individuals of Round 1 exhibited a strong reduction of 4-oxo-octanoic and 4-oxo-decanoic acids (LMMs with ‘diet’ and ‘round’ as fixed variables and ‘colony’ as random variable, FDR-corrected p-values for multiple comparisons; 4-oxo-octanoic acid: untreated *vs.* sucrose-treated, t=15.20, p<0.001, untreated *vs*. tetracycline-treated, t=15.01, p<0.001; 4-oxo-decanoic acid: untreated *vs*. sucrose-treated, t=-13.10, p<0.001, untreated *vs*. tetracycline-treated, t=-13.66, p<0.001). However, changes to these compounds were significant in all pairwise comparisons of all treatment groups, suggesting that it is more related to the isolation than the removal of bacteria (Data set S4). On the other hand, changes in n-C36 and n-C40 were only significant in pairwise comparisons of ants treated with antibiotics and ants reared on their original colonies.

Considering all experimental individuals of Rounds 1 and 2, we found a significant correspondence between chemical profiles (CHC) and qPCR data (RV=0.143, P<0.001; Figure S6). The OTUs *EntAcro1* and *RhiAcro1* co-varied the most with the two metapleural gland acids. There was a significant correspondence between chemical profiles and qPCR gut bacterial abundances also when the acids from the metapleural gland were excluded (RV=0.141, p<0.001). Contrarily, we found no significant correspondences when instead of using the individual CHC and qPCR profiles, we used each aggression pair as a single sample and for input we used the odour differences and ΔΔCt values in aggression pairs (RV=0.122, p=0.707). This suggested that bacterial abundances can explain some of the changes in odour that occurred in the diet treatments (because there is a highly significant correlation between the two data sets), but not to such an extent that they accurately reflect odour differences between individuals that may lead to (a lack of) recognition.

## Discussion

### Experimental treatment affects gut microbial communities

The gut microbial communities of tetracycline-treated ants underwent the most substantial deviation from those of untreated nestmates (two taxa decreased strongly, three slightly and one was not affected; Figure 3), whereas those of sucrose-treated ants exhibited a relatively milder shift (three taxa decreased slightly, two increased slightly and one was not affected). These results show that while the antibiotic suppressed most of the ant microbial community, the sterile sucrose diet affected only mildly the various taxa, altering their relative abundance.

Feeding workers with conspecific faecal droplets only partially restored the original gut microbial communities of sucrose- and tetracycline-treated ants, suggesting that the elimination of strains such as *RhiAcro1* and *EntAcro2* was irreversible. This may imply that some bacteria need to be established during larval development or early adult life and cannot be reintroduced later. In addition, new OTUs emerged (i.e., *EntAcro10*) which may have prevented the original OTUs from tissue re-colonization (cf., 46,47). We also cannot exclude that the tetracycline effect lasted for some time even after its administration was suspended (one week before the preparation of the corresponding microbial DNA samples), potentially interfering with the faecal droplet-mediated microbial gut re-colonization.

Although we provide evidence for which bacterial taxa are most affected by our experimental treatments, amplicon sequencing often does not allow distinguishing bacterial strains with identical 16S sequences (33,48,49). Thus, we cannot exclude the effect of gut bacteria on behaviour to be driven by the interaction of multiple bacterial strains with different metabolic potential, and further research should implement methods that allow considering this.

### Microbiota remodelling affects ant cuticular chemical profiles

Gut bacteria play essential metabolic roles in several ant species, such as the production of metabolites that are absent in host diets or help cover energy needs (50–53), and bacterial removal may thus impair metabolism functions and ant wellbeing. Social withdrawal of unhealthy ant workers has been demonstrated for at least two ant species (54,55), and it is conceivable that the ostracism of the sick ants is driven by changes in CHCs (or other volatile compounds important in communication). We found consistent and significant decreases of two metapleural gland acids (4-oxo-octanoic and 4-oxo-decanoic) (56,57) in sucrose-treated ants, and reductions in these compounds and two linear alkanes (n-c36 and n-c40) in tetracycline-treated individuals. These chemical profile differences may have been caused by symbiotic gut bacteria affecting CHC biosynthesis, either directly by contributing to the CHC pool, or indirectly by affecting host metabolism and therefore host CHC production. This inference is further supported by the fact that CHCs are synthesized in the oenocytes (12), which are heavily colonized by *EntAcro1* bacteria (25) and are the centre of the intermediate metabolism in ants (58).

Tetracycline treatment may not only have affected the gut microbes, but also those living on the ant cuticle (such as the Actinobacteria *Pseudonocardia* (24,34)). Such effects may result from the direct actions of the antibiotic or from indirect effects, such as changes in host metabolism or production of nutrients sustaining cuticular bacterial growth. In *Drosophila*, cuticular microbes have been hypothesized to affect chemical profiles by using CHC as a carbon source or a substrate for degradative enzymes (12), and previous research on *Pogonomyrmex* ants showed that topical antibiotic administration can alter CHC profiles, confirming a possible role of surface microbes in CHC profile determination (18). However, the same kind of antibiotic treatment applied on *Acromyrmex subterraneus* did not affect CHC profiles (19). Work on *Drosophila* suggests that the innate immune response may also mediate CHC profiles (59), and innate immune responses to bacteria affects CHC profiles in honey bees (60). Thus, multiple factors are conceivably involved and likely interact to form worker CHC profiles, and disentangling this complex of host-symbiont interactions and contributions will require extensive further work.

Although we provide correlational evidence for our dietary treatments to act on gut bacteria, which may indirectly affect metapleural gland acids and cuticular hydrocarbons via the metabolism, we cannot exclude other types of effects. Across insect taxa, symbiotic bacteria produce a plethora of volatile semiochemicals (61,62), and studies on *Drosophila* suggest compounds deriving from the metabolism of gut microbes to mediate interactions between individuals (15,63). Eusocial Hymenoptera may have integrated the bacterial metabolism in their chemical-based social dynamics, and ants may thus rely on a social communication system based on chemicals produced by its commensal bacteria. This would allow discriminating against unfamiliar odours from different bacterial communities and would be complementary to the well-studied CHC-mediated nestmate recognition system. Accordingly, we cannot exclude that the altered behaviour observed in our aggression tests may at least partially depend on volatile semiochemicals from bacterial metabolism. In future experiments, nestmate recognition assays should be implemented in which microbiota-remodelled interacting individuals are prevented from relying on non-volatile cuticular social cues.

### Microbiota remodelling affects social interactions

Encounters between antibiotic-treated and untreated nestmates produced the highest aggression levels, suggesting that the antibiotic treatment affects chemical cues that are relevant for social interactions. Aggression due to diet treatment was clearly higher than aggression due to nestmate status and the treatment effect to aggression was directly dependant to the gut bacterial titre: the diet effect on aggression was always clear in sucrose-treated ants but tend to diminish (contrasts were less clear) in tetracycline-treated ants (Figure 2A: nestmate (NM) and non-nestmate (nNM) aggression levels of Tetracycline vs Tetracycline treated ants were almost identical). This further suggested that the gut bacterial communities (or certain taxa) are having an active role in nestmate recognition which possibly disappears when these bacterial symbionts get eliminated.

### The impact of partial restoration of the gut microbiota on social interactions

After faecal droplets administration, nestmate dyadic encounters revealed aggression levels only moderately (and non-significantly) higher than those of Round 1. While this moderate increase in aggression may be an effect of the relatively longer separation of the experimental ant groups (three instead than two weeks in different petri dishes), the lack of significant differences between rounds is consistent with only partial restoration of microbial gut communities. In non-nestmate aggression assays, pairs of tetracycline-treated and tetracycline-vs. sucrose-treated ants again showed low aggression. In contrast, aggression was high in sucrose-treated pairs when interacting individuals were fed with faecal droplets from different colonies. This implies that, although bacterial communities were only partially restored, colony-specific faecal droplets did affect the cues determining behavioural outcomes of the social interactions. This partial restoration also rules out the possibility of changes in behaviour due a direct effect of tetracycline, which is known to impair mitochondrial function and thus potentially could induce confounding effects (64).

## Conclusions

In this study, we seek to take an integrative approach to explore the role of gut microbial symbionts in the social dynamics of a eusocial organism. Remodelling of the ant gut microbiota produced effects on both the chemicals ants display as socially-relevant recognition signals and their resulting behaviours. Our findings suggest that the observed effects mostly depend on two bacterial taxa, the previously identified major gut symbionts of leafcutting ants *EntAcro1* and *RhiAcro1*. Further research will be needed to address the mechanisms underlying the link between these symbionts and behavioural modifications. Either the altered microbial communities may result in chemical profiles that cause individuals to look more like non-nestmates, or microbiota-remodelled ants may be recognized as sick, and aggression towards such individuals could serve to prevent the spread of infections to optimize colony health and efficiency. Regardless, our findings provide evidence that gut bacterial symbionts may be involved in kin recognition by contributing to shape ant CHC signatures, with implications for our understanding of social insect-symbiont evolution.

## References

1. Archie EA, Theis KR. Animal behaviour meets microbial ecology. Anim Behav. 2011 Sep 1;82(3):425–36.

2. Ezenwa VO, Gerardo NM, Inouye DW, Medina M, Xavier JB. Microbiology. Animal behavior and the microbiome. Science. 2012 Oct 12;338(6104):198–9.

3. Dance A. Microbes take charge. Proc Natl Acad Sci U S A. 2014 Feb 11;111(6):2051–3.

4. Cryan JF, Dinan TG. Mind-altering microorganisms: the impact of the gut microbiota on brain and behaviour. Nat Rev Neurosci. 2012 Oct;13(10):701–12.

5. Archie EA, Tung J. Social behavior and the microbiome. Curr Opin Behav Sci. 2015 Dec 1;6(Supplement C):28–34.

6. Clarke G, Grenham S, Scully P, Fitzgerald P, Moloney RD, Shanahan F, et al. The microbiome-gut-brain axis during early life regulates the hippocampal serotonergic system in a sex-dependent manner. Mol Psychiatry. 2013 Jun;18(6):666–73.

7. Heijtz RD, Wang S, Anuar F, Qian Y, Bjorkholm B, Samuelsson A, et al. Normal gut microbiota modulates brain development and behavior. Proc Natl Acad Sci. 2011 Feb 1;108:3047–52.

8. Neufeld KM, Kang N, Bienenstock J, Foster JA. Reduced anxiety-like behavior and central neurochemical change in germ-free mice. Neurogastroenterol Motil Off J Eur Gastrointest Motil Soc. 2011 Mar;23(3):255–64, e119.

9. Buffington SA, Di Prisco GV, Auchtung TA, Ajami NJ, Petrosino JF, Costa- Mattioli M. Microbial Reconstitution Reverses Maternal Diet-Induced Social and Synaptic Deficits in Offspring. Cell. 2016 Jun 16;165(7):1762–75.

10. Leitão-Gonçalves R, Carvalho-Santos Z, Francisco AP, Fioreze GT, Anjos M, Baltazar C, et al. Commensal bacteria and essential amino acids control food choice behavior and reproduction. PLoS Biol. 2017 Apr 25;15(4):e2000862.

11. Sharon G, Segal D, Ringo JM, Hefetz A, Zilber-Rosenberg I, Rosenberg E. Commensal bacteria play a role in mating preference of Drosophila melanogaster. Proc Natl Acad Sci U S A. 2010 Nov 16;107(46):20051–6.

12. Ringo J, Sharon G, Segal D. Bacteria-induced sexual isolation in Drosophila. Fly (Austin). 2011 Dec;5(4):310–5.

13. Arbuthnott D, Levin TC, Promislow DEL. The impacts of Wolbachia and the microbiome on mate choice in Drosophila melanogaster. J Evol Biol. 2016 Feb;29(2):461–8.

14. Morimoto J, Simpson SJ, Ponton F. Direct and trans-generational effects of male and female gut microbiota in Drosophila melanogaster. Biol Lett. 2017 Jul;13(7).

15. Lizé A, McKay R, Lewis Z. Kin recognition in Drosophila: the importance of ecology and gut microbiota. ISME J. 2014 Feb;8(2):469–77.

16. Gadau J, Fewell J. Organization of Insect Societies: From Genome to Sociocomplexity. Harvard University Press; 2009. 638 p.

17. Matsuura K. Nestmate recognition mediated by intestinal bacteria in a termite, Reticulitermes speratus. Oikos. 2001 Jan 1;92(1):20–6.

18. Dosmann A, Bahet N, Gordon DM. Experimental modulation of external microbiome affects nestmate recognition in harvester ants (Pogonomyrmex barbatus). PeerJ. 2016 Jan 25;4:e1566.

19. de Souza DJ, Lenoir A, Kasuya MCM, Ribeiro MMR, Devers S, Couceiro J da C, et al. Ectosymbionts and immunity in the leaf-cutting ant Acromyrmex subterraneus subterraneus. Brain Behav Immun. 2013 Feb 1;28(Supplement C):182–7.

20. José de Souza D, Devers S, Lenoir A. Blochmannia endosymbionts and their host, the ant Camponotus fellah: Cuticular hydrocarbons and melanization. C R Biol. 2011 Oct 1;334(10):737–41.

21. Lester PJ, Sébastien A, Suarez AV, Barbieri RF, Gruber MAM. Symbiotic bacterial communities in ants are modified by invasion pathway bottlenecks and alter host behavior. Ecology. 2017 Mar 1;98(3):861–74.

22. Van Borm S, Billen J, Boomsma JJ. The diversity of microorganisms associated with *Acromyrmex* leafcutter ants. Bmc Evol Biol [Internet]. 2002 May 3;2. Available from: ://000207645900001

23. Sapountzis P, Zhukova M, Hansen LH, Sørensen SJ, Schiøtt M, Boomsma JJ. *Acromyrmex* leaf-cutting ants have simple gut microbiota with nitrogen-fixing potential. Appl Environ Microbiol. 2015 Aug 15;81(16):5527–37.

24. Andersen SB, Hansen LH, Sapountzis P, Sørensen SJ, Boomsma JJ. Specificity and stability of the Acromyrmex–Pseudonocardia symbiosis. Mol Ecol. 2013 Aug 1;22(16):4307–21.

25. Sapountzis P, Zhukova M, Hansen LH, Sørensen SJ, Schiøtt M, Boomsma JJ. Acromyrmex Leaf-Cutting Ants Have Simple Gut Microbiota with Nitrogen-Fixing Potential. Appl Environ Microbiol. 2015 Aug 15;81(16):5527–37.

26. Ottoni EB. EthoLog 2.2: a tool for the transcription and timing of behavior observation sessions. Behav Res Methods Instrum Comput J Psychon Soc Inc. 2000 Aug;32(3):446–9.

27. Bates D, Mächler M, Bolker B, Walker S. Fitting Linear Mixed-Effects Models Using **lme4**. J Stat Softw [Internet]. 2015 [cited 2017 Sep 20];67(1). Available from: http://www.jstatsoft.org/v67/i01/

28. Fox J, Weisberg S. An R Companion to Applied Regression. http://socserv.socsci.mcmaster.ca/jfox/Books/Companion. 2011;

29. multcomp citation info [Internet]. [cited 2017 Sep 29]. Available from: https://cran.r-project.org/web/packages/multcomp/citation.html

30. Zhukova M, Sapountzis P, Schiøtt M, Boomsma JJ. Diversity and Transmission of Gut Bacteria in *Atta* and *Acromyrmex* Leaf-Cutting Ants during Development. Front Microbiol [Internet]. 2017 [cited 2017 Oct 10];8. Available from: https://www.frontiersin.org/articles/10.3389/fmicb.2017.01942/full

31. Ahrens ME, Shoemaker D. Evolutionary history of Wolbachia infections in the fire ant Solenopsis invicta. Bmc Evol Biol. 2005 May 31;5:35.

32. Nygaard S, Zhang G, Schiøtt M, Li C, Wurm Y, Hu H, et al. The genome of the leaf-cutting ant Acromyrmex echinatior suggests key adaptations to advanced social life and fungus farming. Genome Res. 2011 Jan 8;21(8):1339–48.

33. Andersen SB, Boye M, Nash DR, Boomsma JJ. Dynamic *Wolbachia* prevalence in *Acromyrmex* leaf-cutting ants: potential for a nutritional symbiosis. J Evol Biol. 2012 Jul;25(7):1340–50.

34. Andersen SB, Yek SH, Nash DR, Boomsma JJ. Interaction specificity between leaf-cutting ants and vertically transmitted Pseudonocardia bacteria. Bmc Evol Biol. 2015 Feb 25;15:27.

35. Schloss PD, Westcott SL, Ryabin T, Hall JR, Hartmann M, Hollister EB, et al. Introducing mothur: Open-Source, Platform-Independent, Community-Supported Software for Describing and Comparing Microbial Communities. Appl Environ Microbiol. 2009 Jan 12;75(23):7537–41.

36. Pfaffl MW, Horgan GW, Dempfle L. Relative expression software tool (REST©) for group-wise comparison and statistical analysis of relative expression results in real-time PCR. Nucleic Acids Res. 2002 May 1;30(9):e36.

37. Pfaffl MW. Quantification strategies in real-time PCR. A–Z Quant PCR. 2004 Jan 1;89–113.

38. Oksanen J, Blanchet FG, Friendly M, Kindt R, Legendre P, McGlinn D, et al. vegan: Community Ecology Package [Internet]. 2017 [cited 2017 Nov 19]. Available from: https://cran.r-project.org/web/packages/vegan/index.html

39. Fox J, Weisberg S, Friendly M, Hong J, Andersen R, Firth D, et al. effects: Effect Displays for Linear, Generalized Linear, and Other Models [Internet]. 2017 [cited 2017 Nov 19]. Available from: https://cran.r-project.org/web/packages/effects/index.html

40. Hope RM. Rmisc: Rmisc: Ryan Miscellaneous [Internet]. 2013 [cited 2017 Nov 19]. Available from: https://cran.r-project.org/web/packages/Rmisc/index.html

41. Wickham H, Chang W, RStudio. ggplot2: Create Elegant Data Visualisations Using the Grammar of Graphics [Internet]. 2016 [cited 2017 Nov 19]. Available from: https://cran.r-project.org/web/packages/ggplot2/index.html

42. Caporaso JG, Lauber CL, Walters WA, Berg-Lyons D, Huntley J, Fierer N, et al. Ultra-high-throughput microbial community analysis on the Illumina HiSeq and MiSeq platforms. ISME J. 2012 Aug;6(8):1621–4.

43. Aitchison J (John). The statistical analysis of compositional data [Internet]. Chapman and Hall; 1986 [cited 2017 Sep 20]. Available from: http://www.bcin.ca/Interface/openbcin.cgi?submit=submit&Chinkey=123671

44. Hothorn T, Bretz F, Westfall P. Simultaneous Inference in General Parametric Models. Biom J. 2008 Jun 1;50(3):346–63.

45. Dolédec S, Chessel D. Co-inertia analysis: an alternative method for studying species–environment relationships. Freshw Biol. 1994 Jun 1;31(3):277–94.

46. Salem H, Florez L, Gerardo N, Kaltenpoth M. An out-of-body experience: the extracellular dimension for the transmission of mutualistic bacteria in insects. Proc R Soc B Biol Sci [Internet]. 2015 Apr 7 [cited 2016 Jul 25];282(1804). Available from: http://www.ncbi.nlm.nih.gov/pmc/articles/PMC4375872/

47. Anderson KE, Rodrigues PAP, Mott BM, Maes P, Corby-Harris V. Ecological Succession in the Honey Bee Gut: Shift in Lactobacillus Strain Dominance During Early Adult Development. Microb Ecol. 2015 Dec 19;71(4):1008–19.

48. Engel P, Stepanauskas R, Moran NA. Hidden diversity in honey bee gut symbionts detected by single-cell genomics. PLoS Genet. 2014 Sep;10(9):e1004596.

49. Kuo C-H. Scrambled and not-so-tiny genomes of fungal endosymbionts. Proc Natl Acad Sci U S A. 2015 Jun 23;112(25):7622–3.

50. Russell JA, Moreau CS, Goldman-Huertas B, Fujiwara M, Lohman DJ, Pierce NE. Bacterial gut symbionts are tightly linked with the evolution of herbivory in ants. Proc Natl Acad Sci. 2009 Dec 15;106(50):21236–41.

51. de Souza DJ, Bezier A, Depoix D, Drezen JM, Lenoir A. Blochmannia endosymbionts improve colony growth and immune defence in the ant Camponotus fellah. Bmc Microbiol. 2009;9:29.

52. Feldhaar H, Straka J, Krischke M, Berthold K, Stoll S, Mueller MJ, et al. Nutritional upgrading for omnivorous carpenter ants by the endosymbiont *Blochmannia*. Bmc Biol. 2007;5:48.

53. Hu Y, Sanders JG, Łukasik P, D’Amelio CL, Millar JS, Vann DR, et al. Nitrogen conservation, conserved: 46 million years of N-recycling by the core symbionts of turtle ants. bioRxiv. 2017 Sep 7;185314.

54. Heinze J, Walter B. Moribund Ants Leave Their Nests to Die in Social Isolation. Curr Biol. 2010 Feb 9;20(3):249–52.

55. Bos N, Lefèvre T, Jensen AB, D’ettorre P. Sick ants become unsociable. J Evol Biol. 2012 Feb 1;25(2):342–51.

56. Ortius-Lechner D, Maile R, Morgan ED, Boomsma JJ. Metapleural Gland Secretion of the Leaf-cutter Ant Acromyrmex octospinosus: New Compounds and Their Functional Significance. J Chem Ecol. 2000 Jul 1;26(7):1667–83.

57. Larsen J, Fouks B, Bos N, d’Ettorre P, Nehring V. Variation in nestmate recognition ability among polymorphic leaf-cutting ant workers. J Insect Physiol. 2014 Nov 1;70:59–66.

58. Arrese EL, Soulages JL. Insect fat body: energy, metabolism, and regulation. Annu Rev Entomol. 2010;55:207–25.

59. Ryu J-H, Ha E-M, Lee W-J. Innate immunity and gut–microbe mutualism in Drosophila. Dev Comp Immunol. 2010 Apr 1;34(4):369–76.

60. Richard F-J, Aubert A, Grozinger CM. Modulation of social interactions by immune stimulation in honey bee, Apis mellifera, workers. Bmc Biol. 2008 Nov 17;6:50.

61. Davis TS, Crippen TL, Hofstetter RW, Tomberlin JK. Microbial volatile emissions as insect semiochemicals. J Chem Ecol. 2013 Jul;39(7):840–59.

62. Ezenwa VO, Williams AE. Microbes and animal olfactory communication: Where do we go from here? BioEssays News Rev Mol Cell Dev Biol. 2014 Sep;36(9):847–54.

63. Venu I, Durisko Z, Xu J, Dukas R. Social attraction mediated by fruit flies’ microbiome. J Exp Biol. 2014 Apr 15;217(8):1346–52.

64. Moullan N, Mouchiroud L, Wang X, Ryu D, Williams EG, Mottis A, et al. Tetracyclines Disturb Mitochondrial Function across Eukaryotic Models: A Call for Caution in Biomedical Research. Cell Rep. 2015 Mar 10;

